# Atlastins mediate selective autophagy of the endoplasmic reticulum

**DOI:** 10.1101/274530

**Authors:** Jin Rui Liang, Emily Lingeman, Saba Ahmed, Jacob Corn

## Abstract

The selective lysosomal degradation (autophagy) of entire organelles is required for cellular homeostasis, and its dysregulation is involved in degenerative disorders such as Parkinson’s Disease. While autophagy of mitochondria (mitophagy) is becoming better understood, other forms of organelle autophagy are relatively unexplored. Here we develope multiple quantitative assays to measure organelle autophagy using flow cytometry, microscopy, and Western blotting. Focusing on autophagy of the endoplasmic reticulum (ER-phagy), we show that these assays allow facile measurement of ER-phagy, and that ER-phagy is inhibited by knockdown of either core autophagy components or the recently reported FAM134B ER-phagy receptor. Using these assays, we further identify that Atlastins, the ER-resident GTPases involved in ER membrane morphology, are key positive effectors of ER-phagy. Atlastin-depleted cells have decreased ER-phagy under starvation conditions, and Atlastin’s role in ER-phagy requires both a functional GTPase domain and proper ER localization. The three Atlastin family members functionally compensate for one another during ER-phagy and may form heteromeric complexes with one another. We also find that Atlastins act downstream of the FAM134B ER-phagy receptor. We propose that during ER-phagy, Atlastins remodel ER membrane to separate pieces of FAM134B-marked ER for efficient autophagosomal engulfment. Human mutations in Atlastins led to hereditary spastic paraplegia, and our results suggest that this disease may be linked to deficiencies in ER-phagy rather than ER morphology.

## Introduction

The selective autophagy of organelles (organellophagy) constitutes a major part of cellular proteostasis and homeostasis. Dysregulation in organellophagy particularly impacts differentiated cells, such as neurons. The most notable example is mitophagy, whereby loss-of-function mutations of mitophagy proteins such as PARKIN and PINK1 have been linked to neurodegenerative diseases such as Parkinson’s Disease^1^. It is now clear that the endoplasmic reticulum (ER) can also be degraded via ER-specific autophagy (ER-phagy).

The endoplasmic reticulum (ER) is a multifunctional organelle that is the major site for protein and lipid synthesis, as well as the quality control of newly synthesized proteins. To prevent the accumulation of toxic protein aggregates, the ER harbors a well-studied quality control pathway known as *ER*-*A*ssociated *D*egradation (ERAD), in which misfolded ER proteins are extracted for destruction by the proteasome^2^. However, under certain conditions such as starvation, fragments of the ER can also be degraded via ER-phagy^3,4^.

During ER-phagy, fragments of the ER are engulfed in their entirety by autophagosomes and sent for destruction in acidified lysosomes. This process is analogous to the lysosome-mediated degradation of mitochondria (mitophagy). Originally described in yeast, ER-phagy has recently been demonstrated to mediate ER degradation in higher eukaryotic cells^3,6,7^. Two ER-membrane surface proteins with conserved LC3-interacting regions (LIRs), FAM134B and RTN3L, can act as specific autophagy receptors for distinct parts of the ER^8,9^. In addition, SEC62, a component of the ER translocon, and CCPG1, a ER stress-induced ER-phagy receptor, have also been shown to play a role in ER degradation^10,11^. These specialized ER-phagy receptors allow portions of the larger ER network to be shunted to the core autophagy pathways. ER-phagy is therefore connected to bulk autophagy of the cytoplasm, but may have dedicated upstream logic, signals, and mediators that are only beginning to be elucidated. For example, unlike cytoplasm, the ER is composed of a highly interconnected network that presumably must be separated prior to autophagosomal engulfment. It is currently unclear how the ER network is packaged into discrete components for delivery to autophagosomes by ER-phagy receptors.

Dynamin-like GTPases are dimeric, membrane-tethered proteins that are involved in membrane fusion and fission. Depending on their subcellular localization, these GTPases play important roles in regulating the morphology of the targeted organelles^12,13^. For example, Mitofusins (MFNs) and Optic Atrophy 1 (OPA1) are involved in the dynamic fusion and fission of mitochondria. MFNs and OPA1 can also be post-translationally modified and regulated to signal the remodeling of mitochondria during mitophagy^14,15^.

Analogous to MFNs and OPA1, Atlastins (ATLs) are a group of three ER-integral membrane proteins that maintain the ER in a reticular network during normal homeostasis. ATLs contain an N-terminal GTPase domain and two transmembrane helices close to the C-terminus that span the ER membrane, such that both N and C-termini face the cytosol (**Supp Fig. 1A**). Atlastins are required for the formation of three-way junctions that build the characteristic inter-connected, web-like network of the ER. Depletion of ATLs leads to the collapse of the ER network into long threads^16–18^.

Drawing analogy to the roles of MFNs and OPA1 in mitochondrial morphology and mitophagy and reasoning that the ER must be remodeled prior to autophagic engulfment, we hypothesized that ATLs might facilitate ER-phagy. We developed several new assays to measure organelle autophagy, with a focus on ER-phagy. With these assays in hand, we used CRISPR transcriptional inhibition (CRISPRi) to show that ATLs are required for ER-phagy in human cells during nutrient starvation. The three human ATL family members are expressed at different levels in various cell types and are functionally redundant during ER-phagy. In cells that predominantly express ATL2, we find that ER-phagy requires the N-terminal GTPase domain, proper ER localization through the transmembrane domain, and a C-terminal helical tail that is also required for ER membrane remodeling. Overexpression of FAM134B is sufficient to induce ER-phagy and partial loss of ATL2, suggesting that ATL2 could be a remodeling factor for the same ER subdomain marked by FAM134(B)However, removal of ATL2 during FAM134B overexpression abrogates ER-phagy. Our results uncover a new mediator of ER-phagy that we propose is required to remodel and separate the ER components marked for destruction by the FAM134B ER-phagy receptor.

## Results

### Quantitative assays for ER-phagy

One of the obstacles in studying ER-phagy is the paucity of robust markers to monitor the process^3,8,9^. Commonly used ER protein markers such as calnexin (CANX), PDI, CLIMP63, TRAF-α and REEP5 exhibit only a marginal decrease in protein levels during ER-phagy, and it is difficult to use them to establish quantitative measures of ER-phagy. To overcome these limitations, we developed two complementary assays for ER-phagy.

Firstly, we developed a quantitative <u>E</u>R-<u>A</u>utophagy <u>T</u>andem <u>R</u>eporter (EATR) system that fuses a commonly used tandem eGFP-mCherry autophagic reporter to RAMP4, a subunit of the ER translocon complex (Fig. 1A)^19^. EATR is designed such that the fluorescent proteins point towards the cytoplasm and relies on the lowered stability of eGFP relative to mCherry when delivered to the acidic environment of the lysosome. Cytosolic ER (pH∼7) should be fluorescent in both the eGFP and mCherry channels, whereas ER within a lysosomal vesicle (pH<4) would lose eGFP fluorescence but retain mCherry. This system is inspired by the eGFP-mCherry-LC3 reporter system that is commonly used to study bulk autophagy^20^. This system has also previously been used to show that eGFP-mCherry-FAM134B is degraded via the lysosomal pathway^8^. We placed the reporter under the control of a doxycycline-inducible promoter to induce expression in a defined time period, and hence follow a bolus of ER as it transitions from cytoplasmic to lysosomal and is finally degraded (Fig. 1B).

**Figure 1:**
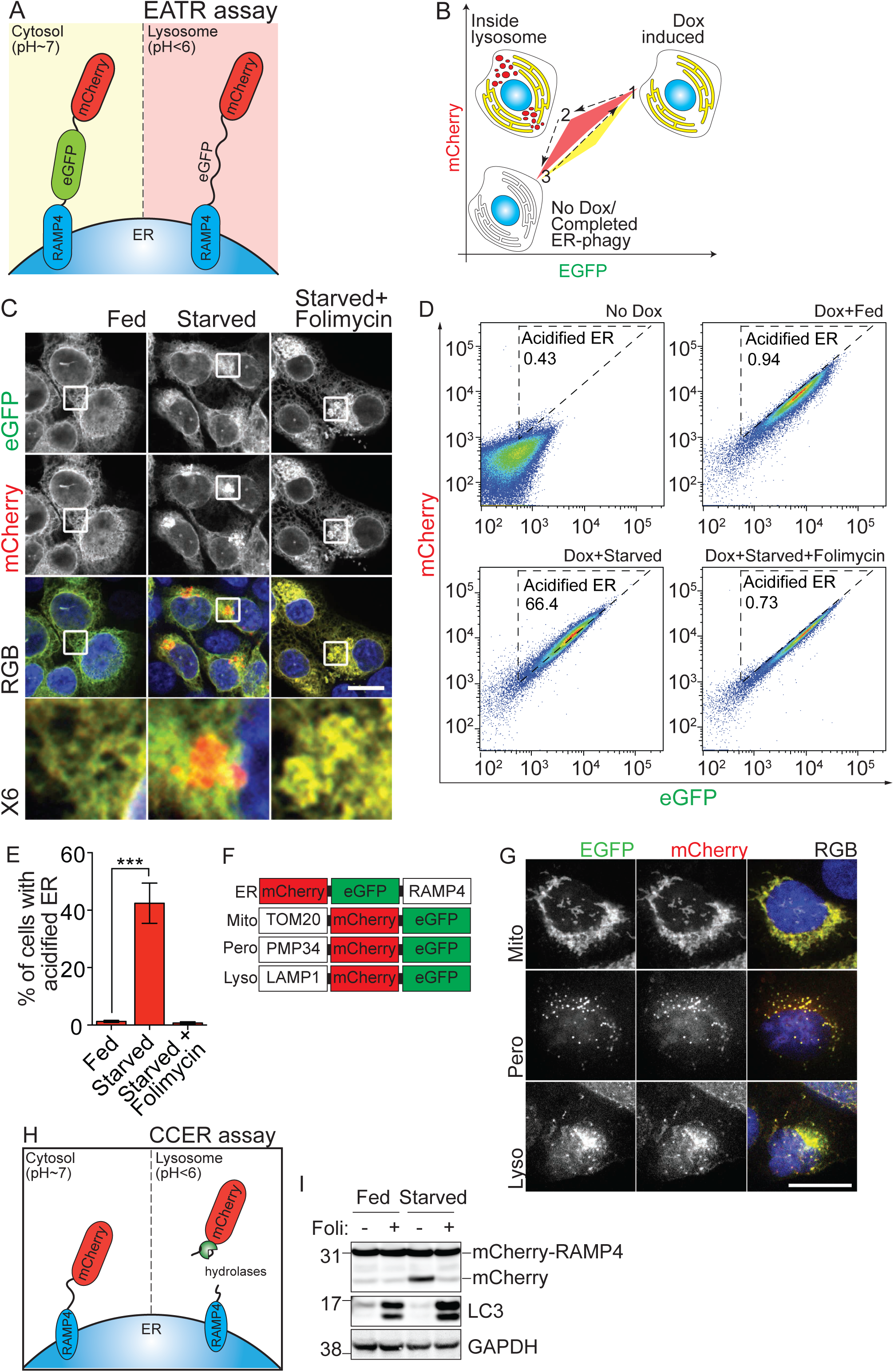
Development of sensitive and quantitative assays to measure ER-phagy. (A).Schematic illustration of the Doxycycline-inducible <u>E</u>R-<u>A</u>utophagy <u>T</u>andem <u>R</u>eporter (EATR) assay featuring a tandem mCherry-eGFP tagged to RAMP4. The environmental pH of cytosolic ER is pH∼7, allowing both eGFP and mCherry to fluoresce normally. During ER-phagy, ER is engulfed and degraded within the lysosome (pH<4), causing the protonation of eGFP and quenching of its fluorescence whereas mCherry protein is relatively stable in the acidic environment. Thus, a switch from GFP^+^/mCherry^+^ to GFP^-^/mCherry^+^ indicates the occurrence of ER-phagy. (B).Schematic of the application of EATR to measure ER-phagy by fluorescence-activated cell sorting (FACS). The progression of ER-phagy is sequentially labeled from 1 to 3. (1) Upon Dox-induction at basal ‘fed’ state, cells express both eGFP and mCherry fluorescence (yellow triangle). (2) During starvation, a fraction of the ER undergoes ER-phagy and loses eGFP fluorescence due to localization within lysosome, resulting in a population shift to the left (red triangle). (3) Complete ER-phagy (with all ER containing the reporter being degraded) will result in complete loss of both eGFP and mCherry expression. Note that complete relocalization of cell population to the bottom left box (complete ER degradation) has yet to be observed as only a small fraction of ER are being degraded during starvation. (C).HCT116 cells expressing the EATR construct show that the reporter localizes to the ER (See also **Supp Fig. 1B**). Starvation induces the formation of perinuclear mCherry-only puncta that colocalizes with the lysosome (**Supp Fig. 1B**). Neutralization of the lysosome using folimycin rescues eGFP fluorescence. Insets represent 6-fold enlargement of the boxed area. Scale bar represents 20µm. (D).HCT116 cells expressing the EATR reporter were doxycycline-induced (4µg/ml) and then starved for 16hrs in EBSS with or without folimycin (100nM). mCherry and eGFP fluorescence was then measured by flow cytometry. The ‘acidified ER’ gate was drawn to exclude the cell population at ‘fed’ condition. Any cell that moves out of the gate represents a decrease in eGFP fluorescence and hence has more ‘acidified ER’, indicative of cells undergoing ER-phagy. Treatment with folimycin to abrogate lysosomal acidification moves all cells out of the ‘acidified ER’ gate. (E).Flow cytometry measurement of ERphagy based on (C) is plotted as percentage of cells with more ‘acidified ER’. Data presented as mean±SD of three biological replicates. P-value indicates two-tailed unpaired T-test with ***P < 0.0005. (F).Schematic illustration of the organellophagy reporters. All reporters with the exception of EATR are fluorescently tagged at the C-terminus to ensure that the fluorescent tag faces the cytosol. (G).U2OS cells expressing each tandem fluorescent reporters with doxycycline regulation. Images were taken at fed state. Scale bar represents 20µm. (H).Schematic illustration of m<u>C</u>herry-<u>C</u>leaved from <u>ER</u> (CCER) assay. ER targeted for degradation results in the lysosomal cleavage of the mCherry tag from RAMP4, resulting in the formation of a smaller, mCherry-only product that can be resolved by Western blotting. (I).HCT116 cells stably expressing CCER were starved for 16hrs in EBSS with or without folimycin (100nM) to measure the accumulation of cleaved mCherry by Western blotting.

We created stable HCT116 cell lines harboring the doxycycline-inducible EATR construct. Using microscopy, we found that EATR is localized to the ER when cells are grown in complete media (“fed” condition). Transitioning reporter cells from “fed” to Earl’s Buffered Saline Solution (EBSS, “starved” condition) induces the formation of mCherry-only ER puncta that are recruited the a perinuclear region, consistent with reports that lysosomes are perinuclear and that autophagic ER cargoes are trafficked to a perinuclear space (Fig. 1C) ^8^,^21^. Treatment with folimycin, an inhibitor of the lysosomal proton pump, rescued GFP fluorescence in perinuclear ER (Fig. 1C). Co-labeling of the ER and lysosome using mCherry-RAMP4 and GFP-LAMP2, respectively, further validated that starvation indeed recruits ER to the lysosome, as evident by the co-localization of mCherry-RAMP4 puncta within GFP-LAMP2 structures (**Supp Fig. 1B**).

We next tested whether EATR could be used to quantify the extent of ER-phagy in a given cell using flow cytometry. Theoretically, relative change in eGFP and mCherry fluorescence intensities should report on delivery of the ER to the acidic lysosomal environment (Fig. 1D). We first validated that doxycycline induction results in double expression of mCherry and eGFP, with each cell having roughly equal amounts of mCherry and eGFP fluorescence. Upon starvation, we observed a decrease in eGFP signal but not mCherry fluorescence, resulting in a shift of cell population into the “acidified ER” gate (Fig. 1D and E). Consistent with our microscopy results, folimycin treatment rescued eGFP fluorescence on the ER, presumably due to de-acidification of the lysosome. Quantifying the number of cells that fall into the acidified ER gate yields a metric of ER-phagy that is reproducible between biological replicates (Fig. 1E). In sum, we find that the EATR system is capable of measuring ER-phagy both visually by microscopy and quantitatively by flow cytometry.

Encouraged by the results with EATR, we adapted the tandem reporter system to study other organellophagies by fusing the tandem fluorescent reporters to different organelle surface proteins, including TOM20 for mitophagy, PMP34 for pexophagy and LAMP1 for lysophagy (Fig. 1F). Similar to the EATR construct, the tandem fluorescent reporters were fused to the organelle-specific protein such that they point towards the cytoplasm and allow for the detection of pH change upon lysosomal localization. In all cases, we observed robust and characteristic localization of all tandem fluorescent reporters to their expected organelles upon doxycycline induction (Fig. 1G).

We further wanted an orthogonal assay for ER-phagy that does not depend upon a change in fluorescence and took inspiration from bulk autophagy assays in yeast that measure lysosomal cleavage of GFP from LC3 as a readout for autophagy^22^. GFP-LC3 is effective in yeast but not human cells, due to differences in lysosomal acidification^19^, but we reasoned that the apparent stability of mCherry in the flow assay could be a replacement for GFP to measure fluorescent tag separation during autophagy. We found that a stably expressed mCherry-RAMP4 fusion (lacking eGFP) is cleaved to free mCherry during starvation, such that Western blotting for mCherry yields two distinct bands (Fig. 1H). Treatment with folimycin prevents release of mCherry, indicating that this separation reports on the internalization of ER by lysosomes (Fig. 1I). For simplicity, we refer to this Western blot assay as m<u>C</u>herry-<u>C</u>leavage from <u>ER</u> (CCER). CCER allows for more sensitive detection of ER-phagy as opposed to measurement of endogenous ER protein degradation, since the appearance and accumulation of the mCherry band from autophagocytosed portion of the ER provides a more definitive readout and more obvious fold change than the marginal loss of endogenous ER proteins relative to the rest of the ER (**Supp Fig. 1C**). Importantly, the timecourse of ER-phagy measured by CCER is consistent with measurements made using endogenous ER resident proteins (**Sup Fig. 1D**).

We further validated that both the EATR and CCER assays are bona fide reporters of ER-phagy by knocking down the FAM134B ER-phagy receptor using short-hairpin RNA^8^. FAM134B depletion led to significant inhibition of ER-phagy as measured by both EATR and CCER (**Fig. 2A-C**). Conversely, we found that overexpression of wild type FAM134B strongly induced ER-phagy in both the EATR and CCER systems even in fed conditions, consistent with the previous reports that FAM134B overexpression is sufficient to trigger ER-phagy (Fig. 2D and E, **Supp Fig. 2A**)^8^. Overexpression of FAM134B with the LIR mutated (FAM134B-LIRmut) did not induce ER-phagy.

**Figure 2:**
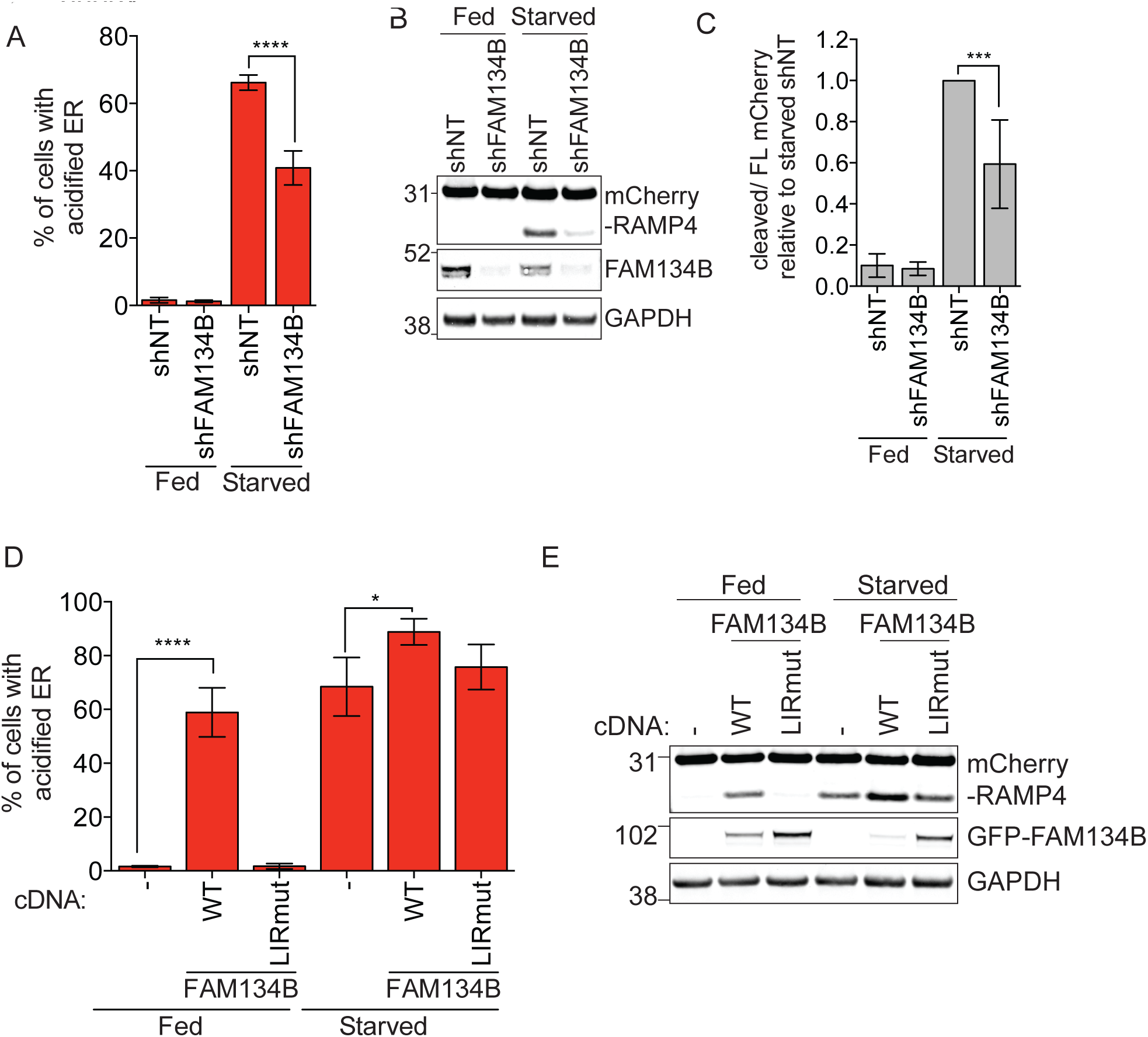
EATR and CCER detect changes in ER-phagy induced by manipulation of a known ER-phagy receptor Er-phagy. (A)EATR HCT116 cells were transfected with short-hairpin RNA targeting FAM134(B)Cells were starved for 16hr prior to FACS measurement. Data presented as mean±SD of three biological replicates. P-value indicates one-way ANOVA with Tukey’s multiple comparisons test. (B)CCER HCT116 cells were treated as in (A), harvested and Western blotted for the indicated proteins. (C)Densitometry measurement of cleaved mCherry versus full-length mCherry-RAMP4 normalized to non-targeting guide RNA (sgNT) at starved condition. Data presented as mean±SD of three biological replicates. P-value indicates one-way ANOVA with Tukey’s multiple comparisons test. (D)EATR HCT116 cells stably expressing FAM134B cDNAs (WT or LIRmut) were starved for 16hrs prior to FACS measurement. Data presented as mean±SD of three biological replicates. P-value indicates one-way ANOVA with Tukey’s multiple comparisons test. (E)CCER HCT116 cells were treated as in (D), harvested and Western blotted for the indicated proteins. A representative blot is shown with quantification from three biological replicates shown in **Supp Fig. 2A**.

### Atlastin knockdown inhibits ER-phagy

Atlastins (ATLs) are ER-resident GTPases involved in ER remodeling, and so represent promising candidates to separate ER for delivery to autophagosomes during ER-phagy. Since Atlastin 1 (ATL1) is preferentially expressed in neuronal cells, we wondered if other Atlastins are differentially expressed in other cell lines^17,23^. Using Western blotting, we found that the levels of each Atlastin vary remarkably in different cellular contexts (**Supp Fig. 3A**). Consistent with previous reports, neuroblastoma SH-SY5Y cells have high level of ATL1 relative to ATL2 and ATL3^17^. However, in other cell lines, ATL1 can be completely absent. Across the panel of cell lines tested, we generally found that there are one or two dominantly expressed Atlastins in each cell line. For example, RPE1 cells only express ATL3 but HepG2 cells only express ATL2. In HCT116 cells, ATL2 is the dominant Atlastin relative to ATL1 and ATL3.

We stably knocked down ATL1, ATL2 and ATL3 via CRISPR-transcriptional inhibition (CRISPRi) in both EATR HCT116-CRISPRi and CCER HCT116-CRISPRi cells that stably express catalytically inactive dCas9 fused with a KRAB transcriptional repressor domain (dCas9-KRAB)^24^. As a control, we knocked down ULK1, a central autophagic regulator kinase required for all forms of autophagy. During starvation, EATR flow cytometry showed that knockdown of ULK1 was sufficient to inhibit ER-phagy, consistent with ER-phagy proceeding through the canonical autophagy pathway (Fig. 3A, **Supp Fig. 3B**).

**Figure 3:**
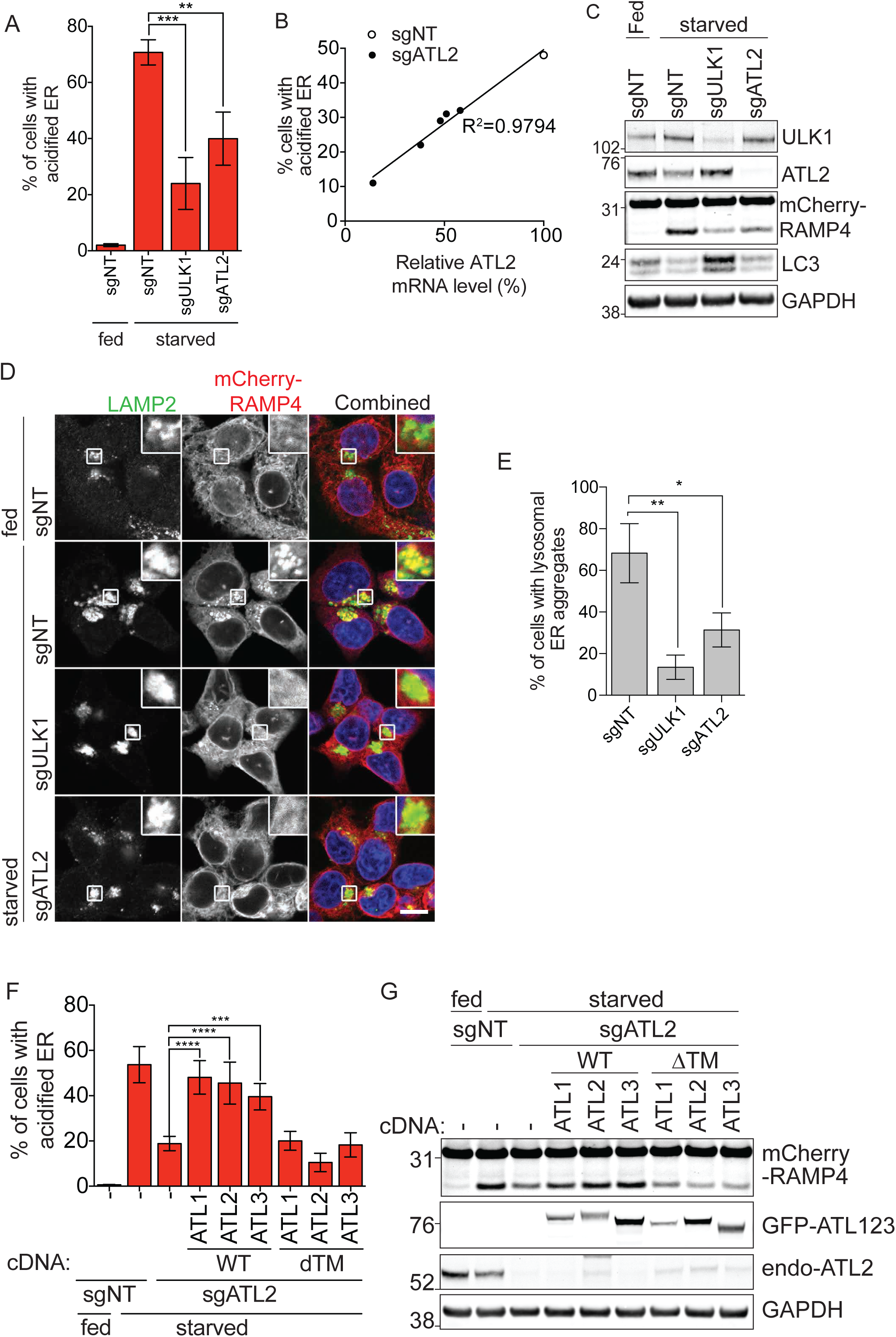
Atlastin knockdown inhibits ER-phagy. (A)EATR CRISPRi HCT116 cells stably expressing sgRNAs targeting either ULK1 or ATL2 were starved for 16hrs and ER-phagy was quantified by flow cytometry. Data presented as mean±SD of three biological replicates. P-value indicates one-way ANOVA with Tukey’s multiple comparisons test. (B)The extent of knockdown from five ATL2-targeting sgRNAs correlates with the extent of ER-phagy inhibition. (C)CCER CRISPRi HCT116 cells stably expressing the sgRNAs shown in (A) were starved for 16hrs, harvested and Western blotted for the indicated proteins. A representative blot is shown with the quantification of three biological replicates shown in **Supp Fig. 3E**. (D)CCER CRISPRi HCT116 cells stably-transduced with the indicated sgRNAs were starved for 16hrs. Cells were fixed and stained with LAMP2 antibody to visualize the lysosomes. Inset represents 3x enlargement of boxed area. Scale bar represents 20µm. (E)Mean cell quantitation (±SD) of three biological replicates from microscopy study shown in (D). An average of 100 cells per condition was quantified. P-value indicates two-tailed unpaired T-test with **P < 0.005 and *P<0.05. (F)EATR CRISPRi HCT116 cells expressing sgATL2 were stably transduced with the indicated GFP-tagged ATL1, 2 or 3 (WT versus ΔTM) and starved for 16hrs. Data presented as mean±SD of three biological replicates. P-value indicates one-way ANOVA with Tukey’s multiple comparisons test. (G)CCER CRISPRi HCT116 cells expressing sgATL2 were stably transduced with the indicated GFP-tagged ATL1, 2 or 3. Cells were starved for 16hrs, harvested and Western blotted for the indicated proteins. A representative blot is shown here with quantification from three biological replicates shown in **Supp Fig. 4B**.

We then used CRISPRi to stably knock down ATL1, ATL2, and ATL3 in hCT116 cells, which express high levels of ATL2. The knockdown efficiency of the sgRNAs for ATL1 and ATL3 were variable, but there was no correlation between sgRNA knockdown efficiency and the extent of ER-phagy (**Supp Fig. 3B-D**). Conversely, we found that depletion of ATL2 inhibited ER-phagy in both EATR and CCER assays, consistent with a role in ER-phagy (Fig. 3A, **Supp Fig. 3C-E**). Moreover, the extent of ER-phagy inhibition correlated very well with the knockdown efficiency of each ATL2 guide RNA (Fig. 3B). We further validated that ATL2 is required for ER-phagy using an exon-targeting ATL2 small interfering RNA (siRNA), and found that siATL2 phenocopies sgATL2’s inhibition of ER-autophagy (**Supp Fig. 3F and G**).

Using CCER, we also observed less accumulation of cleaved mCherry in both ULK1 and ATL2 depleted cells (both sgRNA and siRNA) compared to the non-targeting control cells, consistent with inhibition of ER-phagy observed with EATR (Fig. 3C, **Supp Fig. 3E and G**). However, ULK1 knockdown also resulted in the accumulation of more LC3-I and less conversion into LC3-II, indicating the expected block in general autophagy (Fig 3C, **Supp Fig. 3G**). In contrast, ATL2 depletion by either sgRNA or siRNA did not affect the degradation of LC3, suggesting no inhibition of general autophagy (Fig. 3C, **Supp Fig. 3G**).

We further used confocal microscopy to investigate the effect of ATL2 knockdown on ER-phagy using CCER cells. We observed that sgULK1 and sgATL2 cells show less mCherry-RAMP4 puncta that co-localize with the LAMP2 lysosomal marker (Fig. 3D and E). Combined with robust autophagy progression in ATL2-depleted cells, this suggests that ATL2 depletion prevents targeted ER from autophagic degradation, a role that is fitting with the ER localization of ATL2 and its known roles in the regulation of ER morphology.

### ATL1, ATL2 and ATL3 can functionally compensate for one another during ER-phagy

ATL1, ATL2, and ATL3 have highly similar sequences, with 62-65% overall protein homology and especially at their GTPase domains (**Supp Fig. 4A**). To ask whether the various Atlastins can perform similar functions during ER-phagy, we stably re-expressed cDNAs for ATL1, ATL2 or ATL3 in HCT116 cell lines that are stably knocked down for ATL2. During starvation-induced ER-phagy, re-expression of any one of the three Atlastins was sufficient to rescue ER-phagy (Fig. 3F and G, **Supp Fig. 4B**). By contrast, truncated forms of each Atlastin lacking the ER-localizing transmembrane region (ΔTM) fail to rescue ER-phagy. In addition, immunoprecipitation of ATL2 showed interaction with ATL3 in a manner that is dependent on the GTPase domain but not the TM or C-terminal helix as evident by the lack of interaction with the dimerization defective Arg244Gln (R244Q) mutant but stable interaction with the other constructs (**Supp Fig. 4C**)^17^. This indicates that ATL2 may form a heterodimer with ATL3. Overall, ATL1, ATL2 and ATL3 appear to be functionally interchangeable during ER-phagy and their ER localization is required for this function.

### ATL2 acts downstream of FAM134B during ER-phagy

Previous studies established FAM134B as an ER surface receptor with LC3 interacting regions (LIRs) that recruits autophagic components to the ER. Since ATLs are also localized on the ER membrane, it is possible that they could also play a similar role in autophagosome recruitment. However, while HA-FAM134B co-immunoprecipitates with LC3 in an LIR-dependent manner, ATL2 does not bind to LC3, suggesting that ATL2 serves a different role during ER-phagy (**Supp Fig. 5A**).

Since FAM134B and ATL2 appear to serve distinct functions during ER-phagy, we asked if they act in the same pathway. To this end, we transiently knocked down ATL2 in FAM134B overexpressing cells that upregulate ER-phagy even in the fed state. FAM134B overexpression enhances the degradation of ATL2 under fed condition, and ATL2 protein level is further reduced upon starvation, suggesting that FAM134B is upstream of ATL2 (Fig. 4A). We also found that ATL2 is epistatic to FAM134B overexpression, such that the knockdown of ATL2 reduces FAM134B-induced ER-phagy in both fed and starved conditions (Fig. 4A and B, **Supp Fig. 5B**). Hence, it appears that ATL2 is downstream of FAM134B during ER-phagy.

**Figure 4:**
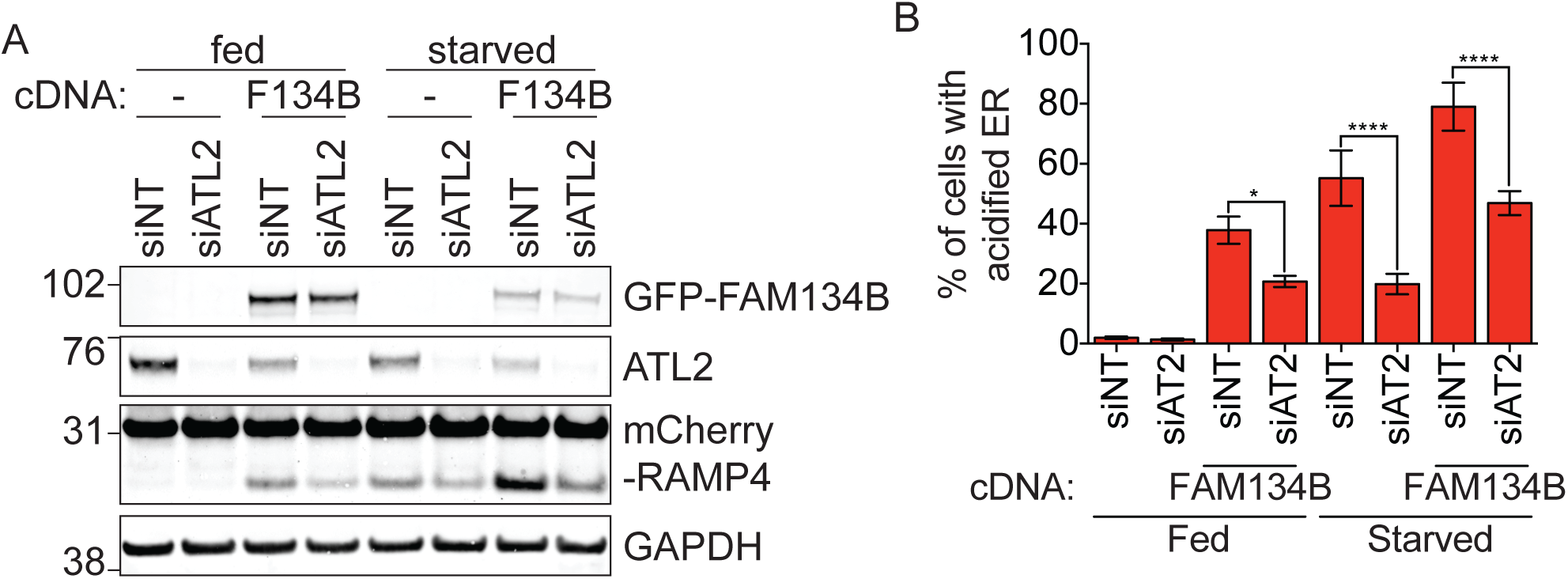
Atlastins act downstream of FAM134B-mediated ER-phagy. (A) CCER CRISPRi HCT116 cells stably expressing GFP-FAM134B and sgATL2 were starved for 16hrs. Representative Western blot is shown here with the fraction of mCherry cleavage measurement from three biological replicates quantified in **Supp Fig. 5B**. (B) EATR CRISPRi HCT116 cells stably expressing GFP-FAM134B and sgATL2 were starved for 16hrs and measured by flow cytometry. Data presented as mean±SD of three biological replicates. P-value indicates one-way ANOVA with Tukey’s multiple comparisons test.

### Atlastin dimerization and GTPase activity are required for ER-phagy

Using ATL2 as an archetypal Atlastin in HCT116 cells, we next asked what molecular features of Atlastins are required for ER-autophagy versus ER morphology. During the maintenance of ER morphology, the various domains of ATL2 perform sequential functions in a GTP-dependent fashion (Fig. 5A)^18^. Upon GTP loading at the GTPase domain, ATL2 is primed for dimerization with another activated ATL2, leading to ER membrane tethering. This is closely followed by ER membrane fusion, a process that requires the C-terminal helix for locaized membrane destabilization{Moss:2011eo}. We generated several constructs of ATL2 to interrogate and compare the importance of each domain for ER morphology and ER-phagy (Fig. 5A).

**Figure 5:**
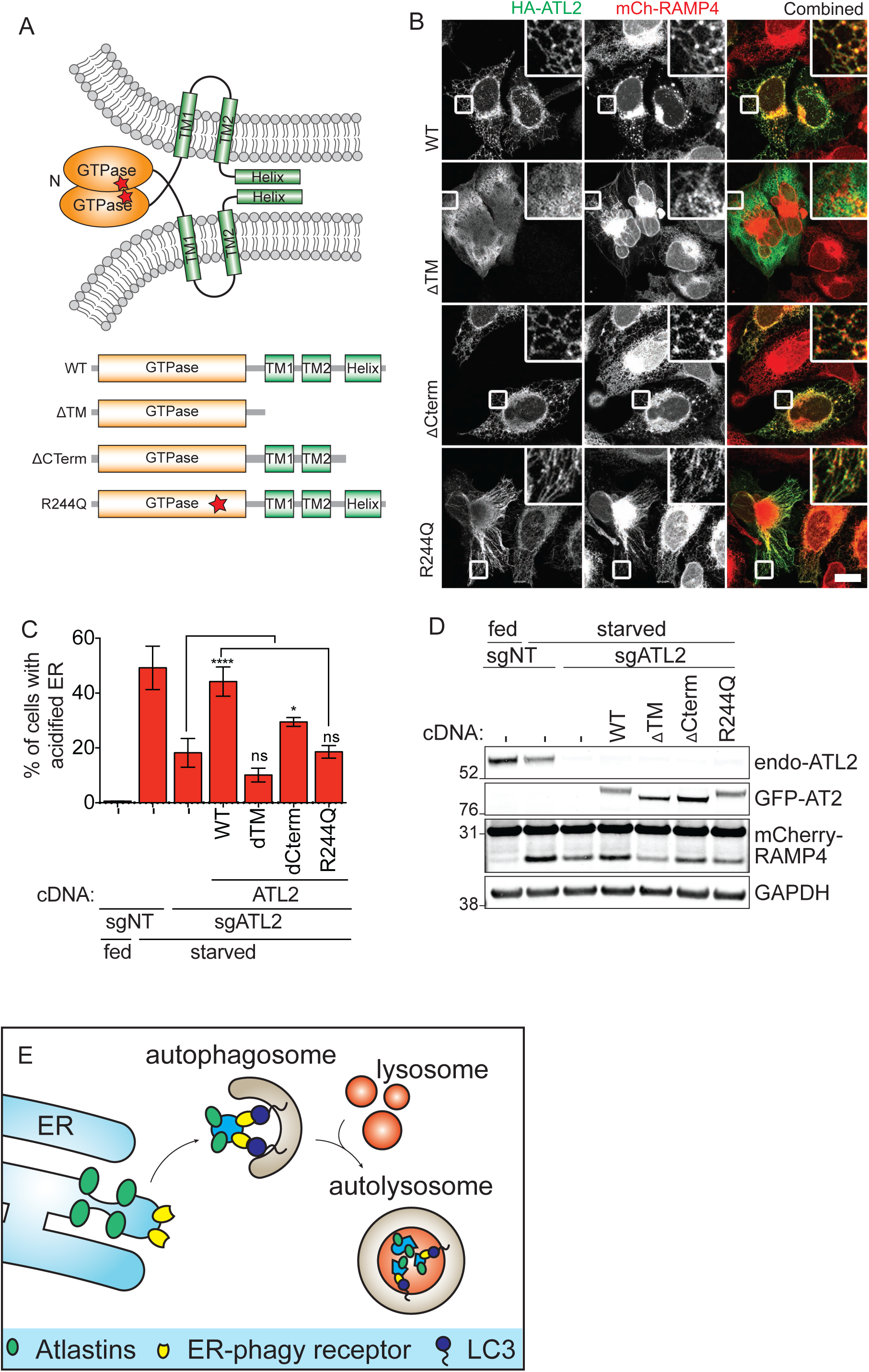
ER-phagy requires a functional Atlastin GTPase domain and proper ER localization. (A) Schematic of the topology and dimerization of Atlastins to facilitate ER membrane fusion. The different truncations and mutations of ATL2 tested shown underneath. (B) U2-OS cells expressing mCherry-RAMP4 were transiently transfected with the indicated HA-tagged mutant versions of ATL2 for 24hrs. Cells were then fixed and imaged to visualize the change in ER morphology. Inset represents 3x enlargement of the boxed areas. Scale bar represents 20µm. (C) EATR CRISPRi HCT116 cells stably expressing sgATL2 and cDNA of ATL2 mutant variants were starved for 16hrs and ER-phagy measured by flow cytometry. Data presented as mean±SD of three biological replicates. P-value indicates one-way ANOVA with Tukey’s multiple comparisons test. (D)CCER CRISPRi HCT116 cells with sgATL2 and stably expressing GFP-tagged version of ATL2 mutants were Western blotted for the indicated proteins. A representative blot is shown here and quantification of three biological replicates shown in **Supp Fig. 6A.** (E) Our data suggests that Atlastins act downstream of FAM134B and we propose that FAM134B acts as the ER-phagy receptor that recruits LC3 to the ER while Atlastins are involved in ER membrane remodeling to deliver FAM134B-marked ER to the autophagosomes.

Overexpressed wild-type ATL2 localizes to the ER network, forming a distinctive reticular network with interspersed puncta at the three-way junctions of the ER (Fig. 5B). During starvation, stable re-expression of wild-type ATL2 restored the diminished ER-phagy caused by depletion of ATL2 (Fig. 5C and D, **Supp Fig. 6A**).

In the presence of endogenous ATL2, overexpressed ATL2 lacking the transmembrane helices (ATL-ΔTM) did not localize to a clearly ER structure. Instead, this construct caused mCherry-RAMP4 marked ER to collapse into an aggregated perinuclear cluster with radiating strands (Fig. 5B). This dominant negative ER morphology phenotype is consistent with previous reports that free ATL2 GTPase domains serve as potent, dosage-dependent inhibitors to endogenous ATL2 by preventing dimerization between endogenous ATL2 molecules and inhibiting ER membrane fusion^18^. We found that re-expression of ATL2-ΔTM during ATL2 knockdown was unable to rescue ER-phagy (Fig. 5C and D, **Supp Fig. 6A**).

We then interrogated the requirement of membrane fusion for ER morphology maintenance and ER-phagy using a truncated form of ATL2 that retains the transmembrane regions but lacks the C-terminal helical domain (ATL2-ΔCterm), We found that overexpression of ATL2-ΔCterm was not dominant negative and had a similar ER morphology to that of wild-type ATL2 at basal state (Fig. 5B). ATL2-ΔCterm expression during ATL2 knockdown partially rescues ER-phagy, indicating that the membrane fusion function of ATL2 is required during ER-phagy (Fig. 5C and D, **Supp Fig. 6A**).

We finally overexpressed an ATL2 mutant that renders the protein incapable of dimerizing (Arg244Gln)^17^. Arg244Gln (R244Q) is equivalent to the human pathogenic Arg217Gln mutation in ATL1 that is implicated in hereditary spastic paraplegia (HSP). Overexpression of Arg244Gln led to ER elongation and lack of reticular connections, but without the collapsed structures observed with the ΔTM construct (Fig. 5B). The ATL2-R244Q mutant was also unable to rescue the loss of ER-phagy induced by ATL2 knockdown. Overall, our data shows that the functional GTPase domain and proper ER localization of ATL2 are required for both ER morphology and ER-phagy regulations (Fig. 5C and D, **Supp Fig. 5A**). Together with our data which indicate that ATL2 is downstream of FAM134B, we propose that FAM134B is the ER-phagy receptor that recruits LC3 to the ER, and Atlastins are responsible for membrane remodeling to deliver FAM134B-marked ER to the autophagy (Fig. 5E).

## Discussion

The mechanisms underlying autophagy of the endoplasmic reticulum are just beginning to be elucidated. We developed two independent systems, EATR and CCER, to report on specific degradation of the ER. The flow cytometry, fluorescence microscopy, and Western blotting assays derived from these systems offer sensitive and quantitative readouts for the progression of ER-phagy. Considering that only a small fraction of ER is actually targeted for ER-phagy at one time, EATR and CCER are more sensitive than measuring endogenous ER protein degradation^3,25^. While EATR provides a quantitative measurement at an individual cell level, CCER can be combined with simultaneous immunoblotting of other ER and autophagy proteins to provide more mechanistic insight into ER-phagy. EATR and CCER therefore serve as complimentary systems to orthogonally explore ER-phagy.

Using EATR and CCER, we found that the ER-resident membrane organizing Atlastin GTPases are required for ER-phagy. While ER-resident autophagy receptors such as FAM134B have been identified, the machinery responsible for isolating ER fragments for delivery to autophagosomes remains unclear^25^. During mitophagy, mitochondria-resident GTPases such as MFNs and MIROs assume a passive role during mitophagy, in the sense that they are ubiquitylated for proteasomal degradation and subsequently lead to mitochondria fragmentation prior to autophagosomal engulfment^15,26^. In contrast, our data suggest that Atlastins play an active role during ER-phagy. Loss of Atlastins represses ER-phagy, and functional Atlastin domains required for ER morphology are also required for ER-phagy. We also found that ATL2 depletion impedes ER-phagy induced by FAM134B overexpression, indicating that ATL2 acts downstream of FAM134B during ER-phagy. Since FAM134B-induced ER-autophagy leads to reductions in ATL2 levels, it also appears that Atlastins act both as an effector and a cargo of FAM134B-mediated ER-phagy.

Our functional studies of ATL2 indicate that a functional GTPase domain, ER localization, and C-terminal helix are all required during ER-phagy. While we cannot conclusively rule out additional roles of Atlastin during starvation, our data suggest that Atlastin’s role during ER-phagy is closely related to its role of ER membrane remodeling. We propose that FAM134B recruits autophagosomes to ER destined for autophagic degradation, and Atlastins locally remodel the FAM134B-marked ER for packaging into autophagosomes (Fig. 5E). ATL1 in particular has known roles in in ER vesiculation during normal homeostasis, and this function may be co-opted during ER-phagy^23^.

Dysregulation in organelle quality control and turnover has detrimental effects on cell fitness and has been linked to multiple diseases. Mounting evidence points towards the importance of organelle quality control for non-regenerative cell types such as neurons. For example, mutations in mitophagy regulators such as PINK1 and Parkin have been reported in patients of Familial Parkinsonism^27^. The FAM134B ER-phagy receptor has likewise been implicated in hereditary sensory neuropathy, a genetic disorder that results in degeneration of sensory nerves of the extremities^28^. Notably, an ATL1 loss-of-function mutation (SPG3A) has been associated with hereditary spastic paraplegia (HSP)^29^. HSP is characterized by the loss of axonal cortical motor neurons, resulting in progressive weakness and spasticity of the lower limbs. ATL1 mutations have previously been proposed to lead to HSP through dysregulation of ER morphology^30^. Considering the similarities in clinical manifestation between FAM134B and ATL1 mutations, combined with our data demonstrating that ATLs act downstream of FAM134B during ER-phagy, we suggest that an inability to execute ER-phagy might be a contributing factor to HSP caused by ATL1 mutations. It remains to be seen whether HSP is concretely caused by deficits in ER-phagy or dysfunctional ER morphology, and further mechanistic studies will hopefully shed light on the molecular basis of the disease.

**Supplementary Figure 1**

(A) Schematic illustrating the topology and dimerization of Atlastins during ER membrane fusion. Both the N-terminal GTPase domain and the C-terminal helix faces the cytosol, with two transmembrane helixes spanning the ER membrane. Atlastins dimerize and undergo GTP hydrolysis to facilitate the fusion of two neighbouring ER membranes.

(B)HCT116 cells stably expressing mCherry-RAMP4 and LAMP2-eGFP were starved for 16hrs with or without folimycin treatment. Cells were fixed and imaged to visualize ER and lysosome co-localization. Inset represents 5x enlargement of boxed area. Scale bar represents 20µm.

(C)CCER HCT116 cells were starved for the indicated duration in EBSS (+/-folimycin) to measure the loss of ER resident proteins and proteins involved in general autophagy by Western blotting.

(D)Densitometry measurement of Western blots in (C). Band intensities were normalized to the highest signal.

**Supplementary Figure 2**

(A) Densitometry measurement from **Fig. 2E** of the ratio of cleaved mCherry band versus full-length mCherry-RAMP4 normalized to the starved GFP-FAM134B-WT sample. Data presented as mean±SD of three biological replicates. P-value indicates one-way ANOVA with Tukey’s multiple comparisons test.

**Supplementary Figure 3**

(A)Cell lysates from different cell lines at ‘fed’ condition were loaded at equal amount to measure the protein expression profiles of ATL1, ATL2 and ATL3.

(B)CRISPRi HCT116 cells were stably transduced with sgRNAs targeting ATL1, 2 or 3. Knockdown efficiencies were assessed by qRT-PCR and normalized to β-actin and non-targeting sgRNA (sgNT). Two previously validated ULK1 sgRNAs were used as positive controls.

(C)sgRNAs with the best knockdown efficiencies based on qRT-PCR from (B) were transduced into EATR CRISPRi HCT116 cells. Data presented as mean±SD of six biological replicates. P-value indicates one-way ANOVA with Tukey’s multiple comparisons test.

(D)CCER CRISPRi HCT116 cells were stably transduced with sgRNA targeting the indicated genes and starved for 16hrs. Cell lysates were harvested and Western blotted for the indicated proteins.

(E)Densitometry measurement from Fig 3C of the ratio of cleaved mCherry band versus full-length mCherry-RAMP4 normalized to the starved NT sample. Data presented as mean±SD of three biological replicates. P-value indicates one-way ANOVA with Tukey’s multiple comparisons test.

(F)EATR CRISPRi HCT116 cells were transiently transfected with siRNA targeting either ULK1 or ATL2 for 48hrs prior to starvation for 16hrs. Data presented as mean±SD of three biological replicates. P-value indicates one-way ANOVA with Tukey’s multiple comparisons test.

(G)CCER CRISPRi HCT116 cells were transiently transfected with siRNA targeting either ULK1 or ATL2 for 48hrs prior to starvation for 16hrs. A representative blot is shown.

**Supplementary Figure 4**

(A)Protein sequence alignment between human ATL1, ATL2 and ATL3 performed using Clustal Omega. The boxed areas indicate the respective GTPase and transmembrane domains.

(B)Densitometry measurement from Fig. 3G of the ratio of cleaved mCherry band versus full-length mCherry-RAMP4 normalized to the starved NT sample. Data presented as mean±SD of three biological replicates. P-value indicates one-way ANOVA with Tukey’s multiple comparisons test.

(C)HCT116 cells stably expressing HA-tagged ATL2 or its mutant variants were immunoprecipitated with anti-HA antibody to measure interaction with ATL3.

**Supplementary Figure 5**

(A)HEK293T cells were transfected with the indicated HA-tagged constructs for 24hrs. Cells were then treated with folimycin for 2hrs prior to HA-immunoprecipitation and Western blotted for the indicated proteins. The LC3 blot was imaged at two different intensities to visualize the input and the HA-IP lanes.

(B)Densitometry measurement from Fig. 4A of the ratio of cleaved mCherry band versus full-length mCherry-RAMP4 normalized to the starved siNT with FAM134B overexpression sample. Data presented as mean±SD of three biological replicates. P-value indicates one-way ANOVA with Tukey’s multiple comparisons test.

**Supplementary Figure 6**

(A) Densitometry measurement from Fig. 5D of the ratio of cleaved mCherry band versus full-length mCherry-RAMP4 normalized to the starved NT sample. Data presented as mean±SD of three biological replicates. P-value indicates one-way ANOVA with Tukey’s multiple comparisons test.

## Materials and Method

### sgRNA, shRNA, siRNA and cDNA plasmid cloning procedures

Information for the sgRNA sequences targeting the transcription start site (TSS) of each gene were obtained from the Weissman CRISPRi-v2 library^1^. In each case, the top 5 guides were cloned into pGL1-library vector (Addgene # 84832) as previously described^1^. Knockdown efficiency for each guide was either validated by qRT-PCR or western blotting. A complete list of all sgRNA constructs used in this study is listed in **Supp Table 1.**

Short hairpin (sh) non-targeting (NT) and shFAM134B were cloned into pLKO.1 puro construct (Addgene #8453) according to protocol available at Addgene website (http://www.addgene.org/tools/protocols/plko/). shRNA sequences used in this study are listed in **Supp Table 1**.

The cDNA of ATL1 was obtained from Harvard PlasmID database (HsCD00326984). cDNAs of ATL2, ATL3 and FAM134B were obtained by PCR amplification of cDNA generated from HCT116 cells. All overexpression constructs were cloned into pLenti-X1-DEST vector (with either neomycin, blastocystin or hygromycin-resistant cassette) via Gibson Assembly^2^. Firstly, a previously cloned pLenti-X1-pEF-mCherry-eGFP-LC3 construct (by LR clonase reaction) was first cut using BamHI and XbaI to liberate the insert. The gene of interest was then Gibson assembled using Gibson Assembly Master Mix (NEB; E2611) together with the desired epitope or fluorescence tag into the pLenti-X1-DEST vector backbone according to manufacturer’s instruction. A complete list of all constructs used in this study is listed in **Supp Table 2.**

### Cell culture

Cell culture was performed at 37°C in a humidified atmosphere containing 5% CO2. HCT116, HEK293T, MCF7, U2-OS, MDA-MB-468, SH-SY5Y, and U2-OS cells were obtained from the Berkeley Cell Culture Facility. All cell lines were cultured in Dulbecco’s Modified Eagle’s medium (DMEM) with high glucose. hTERT-RPE1 cells were cultured in DMEM/F12 with high glucose. All culture media were supplemented with 10% fetal bovine serum (FBS), 100 U/ml penicillin (Gibco), 100 g/ml streptomycin (Gibco), 0.1mM non-essential amino acids and 1 mM sodium pyruvate (Gibco).

### Lentiviral packaging and transduction

Lentiviral packaging of all constructs was performed in HEK293T cells using TransIT®-LT1 Transfection Reagent (Mirus) according to manufacturer’s instructions. Briefly, plasmids were transfected at a ratio of 1:3 (1µg plasmid to 3µl of TransIT-LT1 reagent). The plasmid composition follows 50% plasmid with target of interest 40% ΔVPR plasmid and 10% VSVG plasmid. Lentiviruses were harvested either at 48hr or 72hr post transfection for transduction. Puromycin treatment was performed at 30µg/ml and Neomycin (G418) treatment was performed at 2mg/ml in HCT116 cells. All antibiotic selections were carried out for at least two passages to ensure complete selection.

### Transient Transfection of siRNA and plasmids

Transient transfection of plasmids into HEK293T cells for the purpose of immunoprecipitation was performed using TransIT-LT1 whereas transient transfection of plasmids into U2-OS cells were performed using Lipofectamine 3000 (Invitrogen) according to manufacturer’s instructions at a ratio of 1:3 (1µg plasmid to 3µl of Lipofectamine 3000). Custom ordered siRNA oligoes (Dharmacon) were transiently transfected into targeted cells using RNAiMAX (Invitrogen) according to manufacturer’s instructions. For each well of a 12-well plate, 120pmol of siRNA were diluted in 50µl of OptiMEM. 3.6µl of RNAiMAX was diluted in 50µl of OptiMEM. The diluted siRNA was then added to the diluted RNAiMAX mixture and incubated at room temperature for 5min prior to adding into cell culture. A complete list of all siRNA constructs used in this study is listed in **Supp Table 3.**

### Generation of CRISPRi HCT116 cells

HCT116 cells were transduced with lentivirus with an EF1a-dCas9-HA-BFP-KRAB-NLS construct to generate a pool of CRISPRi HCT116 cells. Cells were FACS sorted and single-cell seeded based on BFP expression. Individual clones were validated by probing for HA tag expression via Western blot. The clones were transduced with a panel of previously validated sgRNAs (SEL1, SYVN, UBE4A, DPH1 and FUT4) using and knockdown efficiency of each clone was assessed by q-RT-PCR. The selected CRISPRi HCT116 clone was further transfected with either TET-On-mCherry-GFP-RAMP4 (for EATR assay) or mCherry-RAMP4 (for CCER assay) and clonal cell lines were generated for uniform doxycycline response and expression of the ER markers. For simplicity, the two cell lines were named EATR CRISPRi HCT116 and CCER CRISPRi HCT116 cells, respectively.

### Cell treatments

For both EATR and CCER assays, cells were seeded and grown for 48hrs prior to starvation. In the case of EATR assay, doxycycline was added at 4µg/ml 24hrs prior to starvation to induce the expression of eGFP-mCherry-RAMP4. Cells were ‘starved’ using Earl’s Buffered Saline Solution (EBSS) with calcium, magnesium and phenol red (Invitrogen, Cat. #24010043) to induce ER-phagy. As opposed to ‘starved’ condition, ‘fed’ condition indicates incubation in normal DMEM with the usual supplements. Unless stated otherwise, co-treatment with folimycin (Milipore) were performed at 100nM final concentration. All starvation and/or drug treatments were carried out for 16hrs unless otherwise indicated.

### Quantitative real-time RT-PCR (qRT-PCR)

Cultured cells were harvested for RNA extraction using Direct-zol RNA miniprep kit (Zymogen) according to manufacturer’s instructions. One microgram of total RNA was used for reverse transcription using Superscript III First Strand Synthesis SuperMix for qRT-PCR (Invitrogen) according to manufacturer’s instructions. qRT-PCR was performed in triplicates using Fast SYBR Green Master Mix (Applied Bioystems) and StepOne Plus Real-Time PCR System (Applied Biosystems). Samples were analyzed with 2-step amplification and melt curves were analysed after 40 cycles. The Ct value for test genes was normalized to β-Actin and the expression of each gene was represented as 2-[ΔΔCt] relative to the non-targeting control. All primers used are listed in **Supp Table 4.**

### Western blotting

After treatment, cells were lysed using RIPA buffer (Milipore) supplemented with Halt Protease inhibitor cocktail (ThermoFisher) and Halt Phosphatase inhibitor cocktail (ThermoFisher). Protein concentration was determined by Bradford Assay (VWR) and NuPAGE LDS Sample Buffer (Invitrogen) was added to the concentration normalized protein samples. Samples were resolved on NU-PAGE Noves Bis-Tris 4-12% gels using MES (2-(*N*-morpholino)ethanesulfonic acid) buffer at 200V for 40min and transferred to 0.4µm nitrocellulose membranes at 1.3A, 25V for 15min using semidry transfer apparatus (BioRad). After protein transfer, membranes were blocked in 5% non-fat dry milk in Tris-buffered saline with 1% Tween-20 (TBS-T) for 30min. Primary antibodies were diluted in 5% BSA in TBS-T and western blots were incubated in primary antibody for either 1hr at room temperature or overnight at 4°C on a rocker. Western blots were then washed twice in TBS-T for 5min each time, followed by secondary antibody incubation using Li-Cor near infrared fluorescence secondary antibodies. All blots were scanned using Li-Cor’s Near-InfraRed fluosrescence Odyssey CLx Imaging System with the exception of any goat primary antibody which was visualized by chemiluminescence using BioRad’s Clarity Western ECL Substrate kit and Chemidoc XRS+ system. A complete list of all primary and secondary antibodies used is listed in **Supp Table 5.** Protein densitometry measurement was performed using Li-Cor’s in-house ImageStudio software.

### Immunofluoresecence

Cells were plated on glass coverslips and were grown for 48hrs prior to any starvation or drug treatment. After treatment, cells were fixed with 4% (w/v) paraformaldehyde (Electron Microscopy Sciences) for 15min. Cells were permeabilized using 0.1% Triton X-100 in PBS for all endogenous protein staining except for LC3 staining. For endogenous LC3 staining, cells were first fixed in 4% paraformaldehyde followed by 100% methanol for 10min. Cells were blocked in 1% BSA in PBS for 20min. Primary antibody staining was performed for 1hr in 1% BSA in PBS, followed by two PBS washes for 5min each. Secondary antibody staining was performed for 30min using Alexa Fluor Dyes (Molecular Probes) in 1% BSA in PBS. Coverslips were mounted onto glass slide using ProLong Gold antifade reagent with or without DAPI for nuclear staining. Images were taken using Zeiss LSM 710 Axio Observer confocal microscope with x63 objective lens. A complete list of all primary and secondary antibodies used is listed in **Supp Table 5.**

### Flow Cytometry Analysis of EATR cells

After treatment, cells were trypsinized and resuspended in complete media without fixation for immediate flow cytometry analysis. All EATR ER-phagy experiments were analysed with live cells because we found that cell permeabilization and fixation reverses eGFP quenching due to loss of lysosomal acidification. On average, 5,000 mCherry and eGFP positive cells were analyzed per each sample. Data were analysed using FlowJo software. Single cells were gated based on eGFP and mCherry fluorescence intensity at fed condition to determine the ‘Acidified ER’ gate. Cells that undergo ER-phagy is defined and quantified based on the shift of cell population into the ‘Acidified ER’ gate due to loss of eGFP fluorescence. *Immunoprecipitation*

Co-immunoprecipitation of LC3 with HA-FAM134B or HA-ATL2 was performed in HEK293T cells by transient transfection with 10µg of plasmid for 24hrs. Co-immunoprecipitation of ATL3 with HA-ATL2 constructs were performed in HCT116 cells stably expressing the different HA-ATL2 mutant constructs. Immunoprecipitation of all HA-cDNA was performed using the Pierce Anti-HA Magnetic Beads Kit (ThermoScientific) according to manufacturer’s instruction. Briefly, cells were lysed using the IP-lysis buffer supplemented with Halt Protease Inhibitor Cocktail (ThermoFisher) and Halt Phosphatase inhibitor cocktail (ThermoFisher). Equal amount of lysates (∼5mg) were used to perform immunoprecipitation using 50µl of HA-magnetic bead slurry. Immunoprecipitation was performed at 4’C for 2hrs. The beads were washed twice using ‘high-salt’ IP lysis buffer (IP lysis buffer supplemented with 500mM NaCl), with 5min incubation on a rotor. The final wash was performed using regular IP lysis buffer. Immunoprecipitated proteins were eluted by boiling the samples at 98’C in 1x NuPAGE(tm) LDS Sample Buffer (thermoFisher) for 5min supplemented with NU-PAGE sample reducing agent at final 1x concentration.

